# Expression of *Plasmodium* Major Facilitator Superfamily Protein in Transporters-Δ *Candida* Identifies a Drug Transporter

**DOI:** 10.1101/2024.02.18.580920

**Authors:** Preeti Maurya, Mohit Kumar, Ravi Jain, Harshita Singh, Haider Thaer Abdulhameed Almuqdadi, Aashima Gupta, Christoph Arenz, Naseem A. Gaur, Shailja Singh

## Abstract

**Aim:** To assess the functional relevance of a putative Major Facilitator Superfamily (MFS) protein (PF3D7_0210300; “*Pf*MFS_DT”) as a drug transporter, using *Candida glabrata* for orthologous protein expression.

**Methods:** CDS encoding *Pf*MFS_DT was integrated into the genome of genetically engineered *C. glabrata* strain MSY8 via homologous recombination, followed by assessing its functional relevance as a drug transporter.

**Results and Conclusion:** The modified *C. glabrata* strain exhibited plasma membrane localization of *Pf*MFS_DT and characteristics of an MFS transporter, conferring resistance to antifungals, ketoconazole and itraconazole. The nanomolar inhibitory effects of the drugs on the intra-erythrocytic growth of *P. falciparum* highlight their antimalarial properties. This study proposes *Pf*MFS_DT as a drug transporter, expanding the repertoire of the currently known antimalarial “resistome”.

## INTRODUCTION

Malaria persists as a significant global health challenge, threatening 40% of the world’s population and an annual toll of half a million lives [1]. Despite concerted research endeavours in fundamental and applied research, the incompatibility of currently available treatment regimens for vulnerable populations, such as children and pregnant women, necessitates the exploration of novel and safe antimalarials. *P. falciparum*, an obligate intracellular parasite, is responsible for the most debilitating forms of malaria, with an ingenious ability to infect and adapt to its host niche, rewiring metabolic capabilities to exploit nutrients for survival. The parasite’s reliance on various transport mechanisms, including those for glucose and amino acids, presents potential drug targets [2].

Within the *Plasmodium*-parasitized erythrocytes, elevated glucose consumption leads to increased formation of monocarboxylates [3]. This dependence of *P. falciparum* on glycolysis for energy production and the maintenance of redox balance highlights metabolic pathways, including the transport of monocarboxylates, as attractive targets for intervention [4,5]. The permeome of *P. falciparum* reveals that 66% of the transporter proteins are essential for the progression of the parasite life cycle [6,7], offering diverse targets for antimalarial development [8,9]. The Major Facilitator Superfamily (MFS), a substantial transporter family in apicomplexan parasites, is implicated with 58 members in related species, such as *Toxoplasma gondii* [10]. MFS transporters play pivotal roles in cellular energy homeostasis and various physiological processes, including nutrient uptake, pH regulation, and the maintenance of intracellular homeostasis [11].

In recent years, there has been growing interest in targeting Monocarboxylate Transporters (MCTs) for therapeutic interventions against diseases spanning cancer, neurological disorders, and infectious diseases [12,13]. A putative *Pf*MFS_DT (PlasmoDB ID: PF3D7_0210300), identified from the permeome of *P. falciparum* [7], underscores its importance in metabolite transportation in the parasite. To identify metabolic bottlenecks and potential drug targets, working with harmonized models as biological mimics, is becoming increasingly important. Since the transporter proteins are less responsive to proteomic tools, their proteomic and functional analysis is cumbersome. Recognizing the challenges, studying these transporters by expressing them in an orthologous system with major drug transporters-deletion genetic background could be a better alternative. Herein, we employed *Candida glabrata* strain MSY8, lacking seven clinically relevant membrane-associated ABC drug transporter genes: *SNQ2*, *AUS1*, *CDR1*, *PDH1*, *YCF1*, *YBT1*, and *YOR1*, along with a gain of function mutation in the *PDR1* transcription factor resulting in a hyper-activation of *CDR1* locus [14]. Since, *C. glabrata* MSY8 is suitable for functional analysis of non-ABC membrane transporters as well [14], we orthologously expressed *Pf*MFS_DT where the masking effects by dominant drug transporters on the activity of other less expressing transporters are reduced due to the deletion of endogenous transporters.

This study aims to evaluate the functional relevance of *Pf*MFS_DT in the transport of drugs, utilizing *C. glabrata* as a yeast model for a heterologous protein expression system, and to further assess its potential as an antimalarial target by identifying small molecule inhibitors. The Complementary Determining Sequence (CDS) encoding *Pf*MFS_DT was codon-optimized for expression in yeast. An integration-based expression plasmid that can integrate *Pf*MFS_DT at the hyperactivated CDR1 locus in the genetically engineered *C. glabrata* strain MSY8 was generated, containing the Nourseothricin N-acetyl Transferase (NAT1) selection marker, CDR1 terminator, promoter regions, YeGFP (at the C-terminus), and unique restriction sites for cloning. *Pf*MFS_DT was precisely directed to integrate at the *Cg*CDR1 locus via homologous recombination. This enabled the expression of fully functional *Pf*MFS_DT on the cell membrane. Recently, Kumari et al. modified *C. glabrata* MSY8 with *Cg*FLR1 (designed as strain ‘MSY13’), an MFS transporter (drug:H + antiporter) that confers resistance to the antifungal 5-flucytosine (5-FC) [14]. MSY13 developed resistance to 5-FC and azoles, including the antifungals: ketoconazole and itraconazole. Similarly, *C. glabrata* strain MSY8 transformed with *Pf*MFS_DT conferred resistance to the antifungals: itraconazole and ketoconazole.

The outcomes of this study have the potential to inform the development of innovative antimalarial therapies, contributing to the ongoing global efforts to combat malaria. Through a combination of bioinformatics, cellular and molecular biology, and pharmacological approaches, we strive to advance our understanding of *P. falciparum* physiology and contribute to the development of urgently needed antimalarial interventions.

## MATERIAL AND METHODS

### Protein architecture and membrane topology

The amino acid sequence of *Pf*MFS_DT (PF3D7_0210300) was retrieved from PlasmoDB, a functional genomic database for *Plasmodium* spp. that provides a resource for data analysis and visualization in a gene-by-gene or genome-wide scale (https://plasmodb.org/plasmo/app/) [15]. Transmembrane helices of *Pf*MFS_DT were predicted using Protter 1.0, the open-source tool for visualization of proteoforms and interactive integration of annotated and predicted sequence features together with experimental proteomic evidence (https://wlab.ethz.ch/protter/start/) [16]. The overall architecture of *Pf*MFS_DT, including transmembrane helices and conserved residues of the Multi-Facilitator Superfamily (MFS) across animalia [17], was illustrated using Illustrator for Biological Sequences (IBS 1.0.3), a tool for visualizing biological sequences (http://ibs.biocuckoo.org/) [18]. Probable amino acid residues of *Pf*MFS_DT corresponding to Na^+^-and lysolipid-binding sites in other eukaryotes were highlighted in green, and those which are conserved across the taxa, animalia, are shown in red (with high consensus value, >90%) and blue (low consensus value, 50%).

### Phylogenetic analysis

Phylogenetic analysis of *Pf*MFS_DT and its orthologs from diverse kingdoms of life, *viz*, archaebacteria, bacteria, apicomplexa, fungi, and animalia was established. To obtain full-length MFS protein sequences from different taxa, BLASTp (https://blast.ncbi.nlm.nih.gov/Blast.cgi) search was performed by using amino acid sequence of *Pf*MFS_DT protein as query sequence, and whole proteomes of Archaebacteria (taxid: 2157), bacteria (taxid:2) apicomplexa (taxid: 5794), fungi (taxid:4751) and animalia (taxid: 33208) as search sets. The maximum likelihood phylogenetic relationship of *Pf*MFS_DT with its orthologs was established using MEGA11, a user-friendly software suite for analyzing DNA and protein sequence data from species and populations (http://www.megasoftware.net/) [19].

### Comparative sequence analysis of *Pf*MFS_DT

Amino acid sequences of MFS proteins from various species, such as, *Danio rerio* (MFSD2A_DR; NCBI: BC085388), *Xenopus tropicalis* (MFSD2A_XT; NCBI: BC123088), *Bos taurus* (MFSD2A_BT; NCBI: BC149727), *Gallus gallus* (MFSD2A_GG; NCBI: XM_417826), *Canis lupus familiaris* (MFSD2A_CLF; NCBI: XP_532546), *Homo sapiens* (MFSD2A_HS; NCBI: NM_032793), *Mus musculus* (MFSD2A_MM; NCBI: NM_029662), *Xenopus tropicalis* (MFSD2B_XT; UniProtKB: A4IH46), *Bos taurus* (MFSD2B_BT; NCBI: XM_010810291), *Gallus gallus* (MFSD2B_GG; NCBI: XM_004935790), *Canis lupus familiaris* (MFSD2B_CLF; NCBI: XM_005630178), *Homo sapiens* (MFSD2B_HS; NCBI: NM_001346880), and *Mus musculus* (MFSD2B_MM; NCBI: NM_001033488) [17] were retrieved from the NCBI database (https://www.ncbi.nlm.nih.gov/). Multiple sequence alignment was performed using the MultAlin tool, which creates sequence (either protein or nucleic acid) alignment from a group of related sequences by using progressive pairwise alignments. (http://multalin.toulouse.inra.fr/multalin/) [20]. Residues with high consensus values (>90%) were denoted in red, while those with low consensus values (>50%, <90%) were denoted in blue. The residues associated with Na^+^- and lysolipid-binding sites were marked with stars, and transmembrane helices are highlighted.

### P. falciparum culture

*P. falciparum* laboratory-adapted strain 3D7 was cultured using standard protocols [21]. Parasites were cultured in RPMI-1640 medium (Gibco^TM^, USA) supplemented with 50 mg/L hypoxanthine (Sigma-Aldrich, USA), 2 gm/L sodium bicarbonate (Sigma-Aldrich, USA), 5 gm/L AlbuMax I, and 10 mg/L Gentamicin (Sigma-Aldrich, USA). Cultures were maintained in 75 cm^2^ (T-75) culture flasks (Corning®, US) using fresh O-positive (O+) human erythrocytes under an ambient mixed gas environment (5% O_2_, 5% CO_2_, and 90% N_2_) at 37°C. Before every experiment, parasite cultures were synchronized with 5% D-sorbitol for two successive intra-erythrocytic proliferative cycles.

### Expression and colocalization of *Pf*MFS_DT in *P. falciparum*

IFAs were conducted on synchronized *P. falciparum* 3D7 cultures to assess *Pf*MFS_DT expression and its co-localization with Merozoite Surface Protein (*Pf*MSP1) during late developmental stages (34 - 46 hpi). Percoll-purified parasitized erythrocytes were isolated, smeared on glass slides, fixed with pre-chilled methanol overnight at −20 °C, and then blocked with 5% BSA (SRL Chemical) in 1X PBS for 1 hour at RT. The slides were probed with anti-*Pf*MFS_DT_Hydrophillic_ mouse serum (1:100) and anti-*Pf*MSP1 rabbit serum (1:300; Sigma-Aldrich) for 1 hour at RT. After washing, the slides were probed with anti-mice Alexa-Fluor 488 (for *Pf*MFS_DT; Thermo Fisher Scientific) and anti-rabbit Alexa fluor 594 (for MSP1; Thermo Fisher Scientific), both at 1:300 dilution and mounted with ProLong Gold antifade reagent with DAPI (4’,6-diamidino-2-phenylindole; Invitrogen). Confocal microscopy (Olympus FLUOVIEW FV3000) was used for image acquisition, and co-localization analysis was performed on a pixel-by-pixel basis using Olympus cellSens Dimension imaging software (https://www.olympus-lifescience.com/en/software/cellsens/).

To assess *Pf*MFS_DT expression in parasites, late-stage parasites (34 - 46 hpi) were isolated through saponification with 5% saponin (Sigma Aldrich). The isolated parasite pellet was lysed using RIPA lysis buffer (30 mM Tris, 150 mM NaCl, 20 mM MgCl_2_, 1mM EDTA, 1 mM β-ME, 0.5% Triton X-100, 1% IGEPAL, 1 mM PMSF, and 0.1% SDS; pH 8.0). The lysate was resolved on 12% SDS-PAGE, transferred to a nitrocellulose membrane (mdi Membrane Technologies), and probed with α-MFS_DT antisera (1:400) for 1 hour at Room Temperature (RT). Subsequently, a secondary α-HRP anti-mice antibody (1:5,000; Sigma-Aldrich) was applied for 1 hour at RT.

### Orthologous model for accessing *Pf*MFS_DT transport activity using *C. glabrata* MSY8

The Complementary Determining Sequence (CDS) encoding *Pf*MFS_DT was codon-optimized for expression in yeast (GenScript Biotech, United States) **(Supplementary Figure S2**; the original and codon-optimized nucleotide sequences of *Pf*MFS-DT are shown in the supplementary material). The *Candida glabrata* BG14-derived MSY8 strain (Δ*Cgsnq2::FRT,* Δ*Cgaus1::FRT,* Δ*Cgcdr1::FRT,* Δ*Cgpdh1::FRT,* Δ*Cgycf1::FRT,* Δ*Cgybt1::FRT,* Δ*Cgyor1::FRT,Cgpdr1::CgPDR1^G840C^ FRT*) [14] was maintained in YPD (1% yeast extract and 2% peptone media supplemented with 2% Dextrose) (Himedia chemicals, India). Yeast cultures were grown at 30 °C and 200 rpm.

For yeast cell transformation, a homologous recombination-based strategy was employed to complement *Pf*MFS_DT, as previously outlined by Kumari et al. in 2020 [14]. In brief, overnight cultures were sub-cultured, grown to an optical density at 600 nm (OD_600_) of 1.5, harvested, washed with sterile water, and resuspended in 100 mM lithium acetate. Following centrifugation at 1,000 xg for 30 seconds, cells were resuspended in a solution containing 50% PEG-3350, Salmon Sperm (SS) DNA (270 µg/mL), 100 mM lithium acetate, and 1 µg of the transformation “cassette.” After 1 hour incubation at 30°C, cells were subjected to a heat shock at 42°C for 15 minutes. Subsequently, the cells were harvested, incubated at 30°C and 160 rpm in YPD medium for 1 hour, harvested again, resuspended in sterile water, plated on Nourseothricin (NAT) selection plates, and then incubated at 30°C for 48 hours.

### Homology modeling of *Pf*MFS_DT

Homology modeling of *Pf*MFS_DT was performed using the Prime application in Schrodinger Maestro Suite 2022-4 (Schrodinger, LLC, New York, NY, 2014). The protein sequence (457 amino acids; accession ID: O96186) was retrieved from the UniProt database [22]. NCBI Protein BLAST against the PDB database identified chain_A of bacterial oxalate transporter OxlT 8HPK_A [23] as the best template for homology modeling, with a sequence similarity of 28.6 % and gaps of 1.5 %. The three-dimensional crystal structure of OxlT 8HPK_A was retrieved from the RCSB PDB database for use as the template.

Amino acids from Phe-32 to Ser-36, Gln-164 to Tyr-188, and Val-240 to Gln-272 of *Pf*MFS_DT were excluded from the modelling due to a lack of corresponding template coordinates. Homology modeling was performed using the Knowledge-based model-building method in Prime, generating one model. The model was refined through loop refinement and energy minimization using the OPLS4 force field in Schrodinger Prime [24]. The minimized model was validated using a Ramachandran plot generated from PROCHECK at the SAVES version 4 server (Structure Analysis and Verification Server; http://services.mbi.ucla.edu/SAVES/) [25]. This model was optimized prior to the docking analysis using the Protein Preparation workflow in Schrodinger Maestro Suite 2022-4 (Schrodinger, LLC, New York, NY, 2014) [26–28].

### *In silico* interaction analysis

#### Protein and Ligand Preparation

Protein preparation for the generated model was conducted using the Protein Preparation Wizard within Schrödinger Maestro 2022-4, optimizing the protein structure’s energy with the OPLS4 force field (Schrodinger, LLC, NY, USA, 2009) [26,27].

Ligands were prepared using the LigPrep module within Schrodinger Maestro 2022-4 (Schrodinger, LLC, NY, USA, 2009), converting 2D structures to 3D structures, optimizing geometry, adjusting chirality and desalting. Ionization and tautomeric states were determined using the Epik module, accommodating pH values ranging from 5 to 9. The libraries underwent minimization using the Optimized Potentials for Liquid Simulations-4 (OPLS-4) force field within Schrodinger software (Li et al., 2011). Subsequently, a single energetically favourable conformation for each ligand was generated for docking analysis [27].

#### Validation of binding site and grid box generation

Binding site validation was performed using the SiteMap tool in Maestro 2022-4 of the Schrodinger suite. The binding site with the highest score was selected for grid generation. Receptor grid boxes were generated at the protein’s active site using the “Glide’s Receptor Grid Generation” module.

#### Glide ligand docking

Ligands (ketoconazole and itraconazole) were docked against the active site of *Pf*MFS_DT using Glide [29] and OPLS-4 [24] force fields. The docking algorithm assessed hydrogen-bonding, hydrophobic, and electrostatic interactions while avoiding steric clashes. The Glide Score scoring function was applied to re-score the obtained poses, and docking was performed with Extra Precision (XP) docking [29–31].

#### Calculation of Binding Free Energy Using Prime/MM-GBSA Approach

Binding free energies of the ligand-receptor complexes were computed using the Prime module from the Schrödinger suite 2022-4. The Molecular Mechanics-Generalized Born Surface Area (MM-GBSA) method, utilizing the OPLS4 force field, VSGB solvent model, and search algorithms, was used [24].

### *Pf*MFS_DT expression in MSY8 strain and assessment of transported molecules

The sensitivity of *Pf*MFS_DT-expressing MSY8 strain (MSY8-*Pf*MFS_DT1) towards antifungal compounds: ketoconazole and itraconazole, was evaluated. Towards this, at the MIC_80_ concentration of the compounds, serial dilution spot assay was performed in the presence of itraconazole (0.031 µg/mL) and ketoconazole (0.002 µg/mL) at their MIC_80_ on MSY8 [14]. Transformed and untransformed yeast cells were grown overnight in YPD medium, suspended in sterile Phosphate-Buffered Saline (PBS) to an OD_600_ of 0.1, spotted on YPD agar plates with the given concentrations of the compounds, and growth differences were recorded after 48 h.

### Ketoconazole and itraconazole inhibit intra-erythrocytic parasite growth

To further investigate the antimalarial properties of ketoconazole and itraconazole, their inhibitory activity was assessed on *P. falciparum* strain 3D7. Parasite culture synchronized at ring stage, with 0.8 - 1% parasitaemia and 2% haematocrit were treated with different concentrations (50, 25, 12.5, 6.25, 3.125, 1.56, 0.78, and 0.39 µM) of the compounds for 72 hours at 37 °C. Giemsa-stained smears were prepared, and parasite growth was determined by counting infected RBCs relative to total erythrocytes using a light microscope.

## RESULTS

### Architecture, phylogeny, and functional residues of *Pf*MFS_DT

Probable amino acid residues involved in Na^+^- and lysolipid-binding were identified and highlighted in the overall architecture of *Pf*MFS_DT **(Figure 1A)**. Residues with high consensus value (>90%; red) and those with low consensus value (50%; blue) are also shown. The helical regions in *Pf*MFS_DT are delineated from H1 to H12. Topology prediction revealed the presence of 12 transmembrane helices, with both N- and C-termini of *Pf*MFS_DT situated intracellularly **(Figure 1B)**. The maximum likelihood phylogenetic analysis demonstrated a convergent evolutionary pathway of *Pf*MFS_DT with MFS proteins from the related apicomplexan, *Toxoplasma gondii* **(Figure 1C)**. Branches with red, blue, and orange filled circles represent MFS proteins from *Plasmodium*, *Toxoplasma*, and Theileria sp., respectively. Further detailed evolutionary relationship of *Pf*MFS_DT with its orthologs from various taxa (archaebacteria, bacteria, apicomplexa, fungi, and animalia) is shown in **Supplementary Figure S1**. The MultAlin-based sequence alignment with well-characterized MFS orthologs in animals illustrated conservation patterns and correlations in *Pf*MFS_DT **(Figure 1D)**. Amino acid residues with high consensus value (>90%) are shaded in red and those with low consensus value (50%) are shaded in blue. Twelve transmembrane helices are also indicated. Probable amino acid residues of *Pf*MFS_DT corresponding to Na+- and lysolipid-binding sites in other eukaryotes were highlighted with stars.

**Figure 1:**
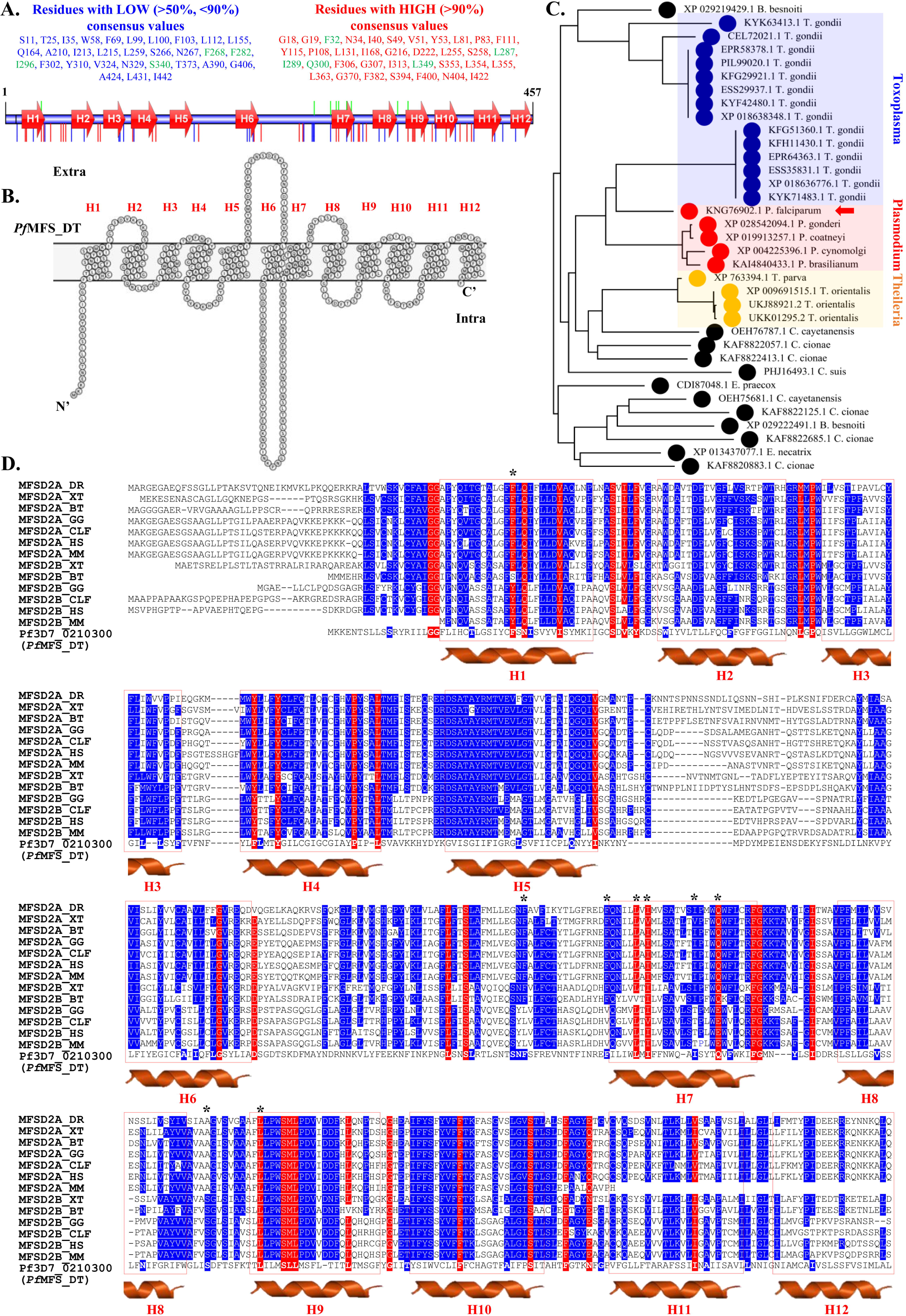
**(A) Overall *Pf*MFS_DT architecture** as deduced using IBS 1.0.3 [18]. Probable amino acid residues involved in Na^+^- and lysolipid-binding, along with transmembrane helices (H1 to H12) are shown. **(B)** *Pf*MFS_DT topology prediction using Protter 1.0 [16]. **(C) Maximum likelihood phylogenetic analysis** generated using MEGA11 [19] demonstrates a convergent evolutionary pathway of *Pf*MFS_DT with MFS proteins from *Toxoplasma gondii*. Red, blue, and orange filled circles represent MFS proteins from *Plasmodium*, *Toxoplasma*, and Theileria sp., respectively. **(D)** MultAlin-based sequence alignment of *Pf*MFS_DT with well-characterized orthologs in animal species illustrated conservation patterns and correlations. Multiple sequence alignment was performed using the MultAlin tool [20]. Red: residues with high consensus values (>90%); Blue: residues with low consensus values (>50%, <90%); and, Stars: residues associated with Na^+^- and lysolipid-binding.

### *Pf*MFS_DT is expressed by *P. falciparum* and localizes on the cell membrane

IFAs were conducted on synchronized *P. falciparum* 3D7 cultures to assess *Pf*MFS_DT expression and its co-localization with *Pf*MSP1 during late developmental stages (34 - 46 hpi). *Pf*MFS_DT expression was observed on the plasma membrane of late-stage trophozoites and schizonts of the *P. falciparum* 3D7 strain, co-localizing with *Pf*MSP1 **(Figure 2A)**, which was confirmed by Western blot analysis revealing the expression of *Pf*MFS_DT (51.8 kD) during the intra-erythrocytic stages of *P. falciparum* **(Figure 2B)**.

**Figure 2:**
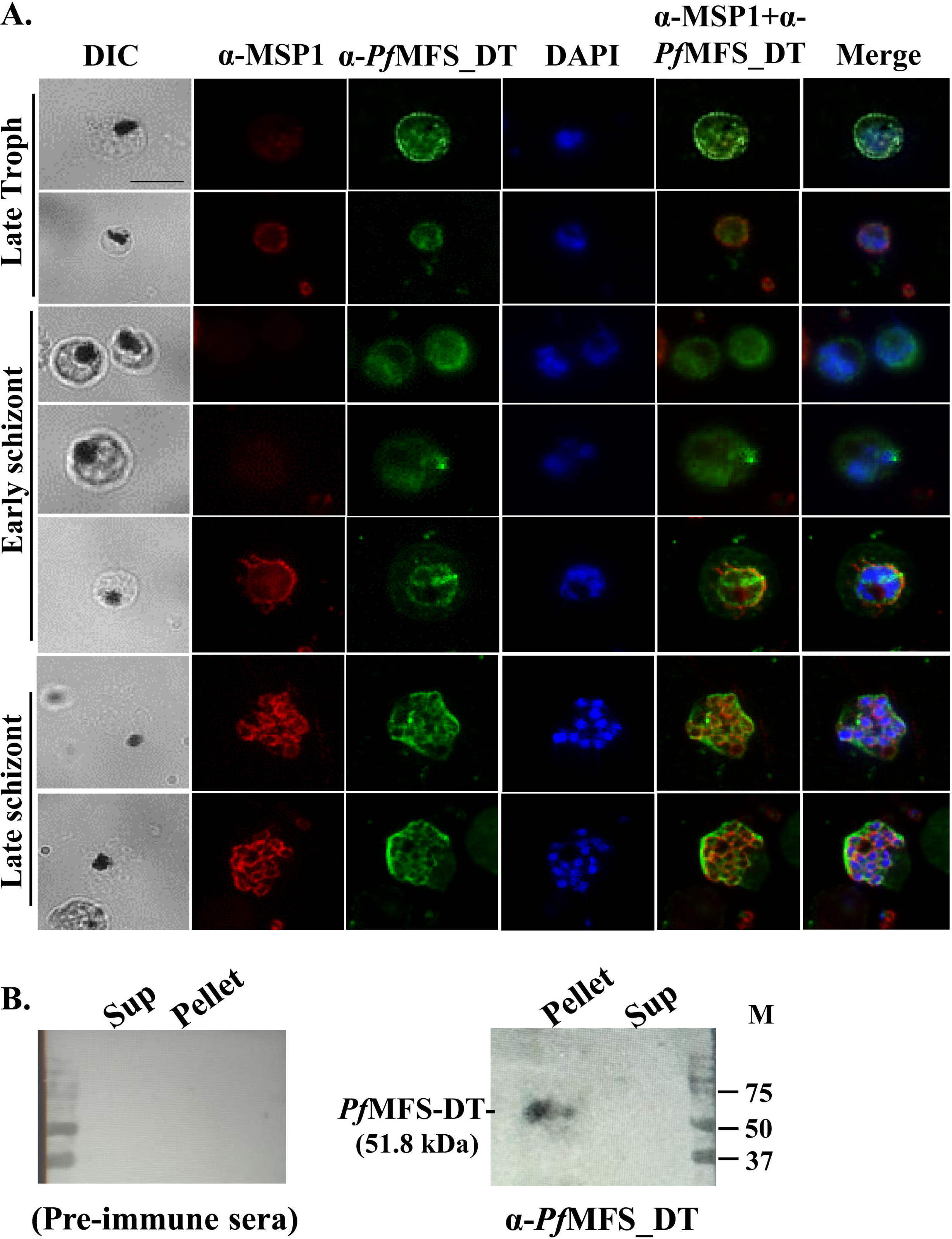
Expression and co-localization of *Pf*MFS_DT with *Pf*MSP1. **(A)** IFA to check the localization of *Pf*MFS_DT in the parasite using anti-*Pf*MFS_DT_Hydrophillic_ mouse serum; localization of MSP1 served as positive control. *Pf*MFS_DT localizes towards the parasite cell membrane and co-localizes with *Pf*MSP1. **(B)** Western blotting confirming *Pf*MFS_DT expression during the schizont stage using anti-*Pf*MFS_DT_Hydrophilic_ mouse serum.

### Generation of yeast model: Orthologous expression of *Pf*MFS_DT in ABC transporters-deficient *C. glabrata*

We orthologously expressed *Pf*MFS_DT in the *C. glabrata* MSY8 strain (“MSY8-*Pf*MFS_DT1”) generated by Sonam Kumari *et al.* in 2020 [14], which has disruptions in seven clinically relevant membrane-localized ABC drug transporter genes: *CgSNQ2*, *CgAUS1*, *CgCDR1*, *CgPDH1*, *CgYCF1*, *CgYBT1*, and *CgYOR1*, along with a gain-of-function mutation in the *CgPDR1* transcription factor, resulting in the hyperactivation of the *CgCDR1* locus [14]. *Pf*MFS_DT was specifically directed to integrate at the *CgCDR1* locus to enhance its expression through homologous recombination **(Figure 3A)**. MSY8-*Pf*MFS_DT1 exhibited plasma membrane localization of *Pf*MFS_DT, as evidenced by the detection of GFP through fluorescence microscopy **(Figure 3B)**. This was confirmed by Western blotting and probing *Pf*MFS_DT-GFP with mouse anti-*Pf*MFS_DT_hydrophilic_ (1:400), followed by using anti-mice (1:5,000; Sigma Aldrich) HRP-conjugated secondary antibodies **(Figure 3C)**.

**Figure 3:**
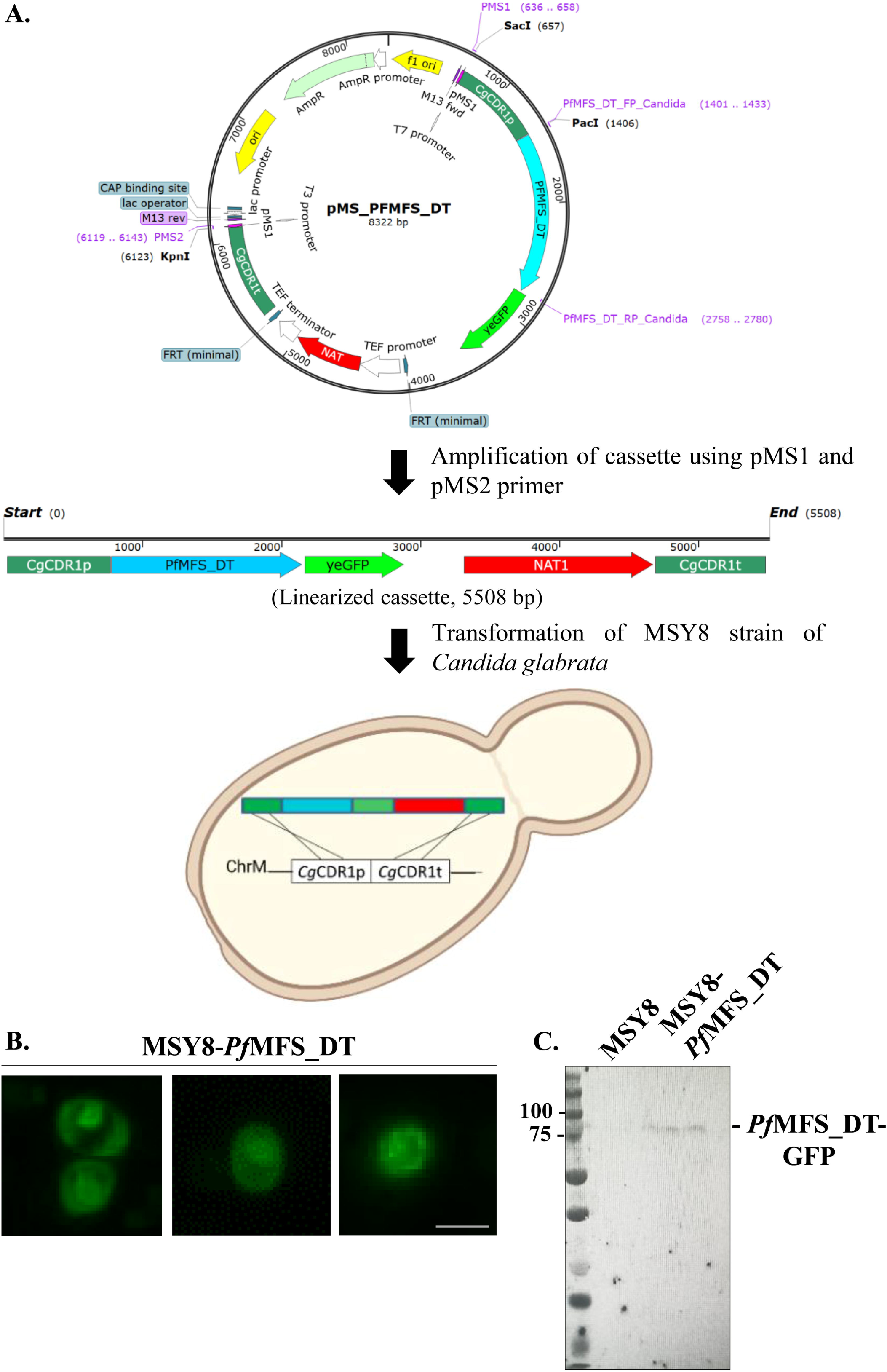
(A) Schematic representation of *Pf*MFS_DT cloning in the pMS vector and the subsequent transformation of the *C. glabrata* strain MSY8. **(B)** Visualization of *Pf*MFS_DT-GFP in transformed *C. glabrata* strain MSY8 strain using fluorescence microscopy, revealing localization towards the cell membrane and nucleus. **(C)** Western blot using anti-*Pf*MFS_DT_Hydrophillic_ mouse serum confirming expression in transformed *C. glabrata* strain MSY8. A desired protein band of ∼79 kDa was observed.

### *Pf*MFS_DT desensitizes *C. glabrata* against the antifungals, ketoconazole and itraconazole

The sensitivity of MSY8-*Pf*MFS_DT1 towards antifungal compounds: ketoconazole and itraconazole was evaluated, which showed rescued growth in the presence of ketoconazole and itraconazole, suggesting the involvement of *Pf*MFS_DT in their transport **(Figure 4)**.

**Figure 4:**
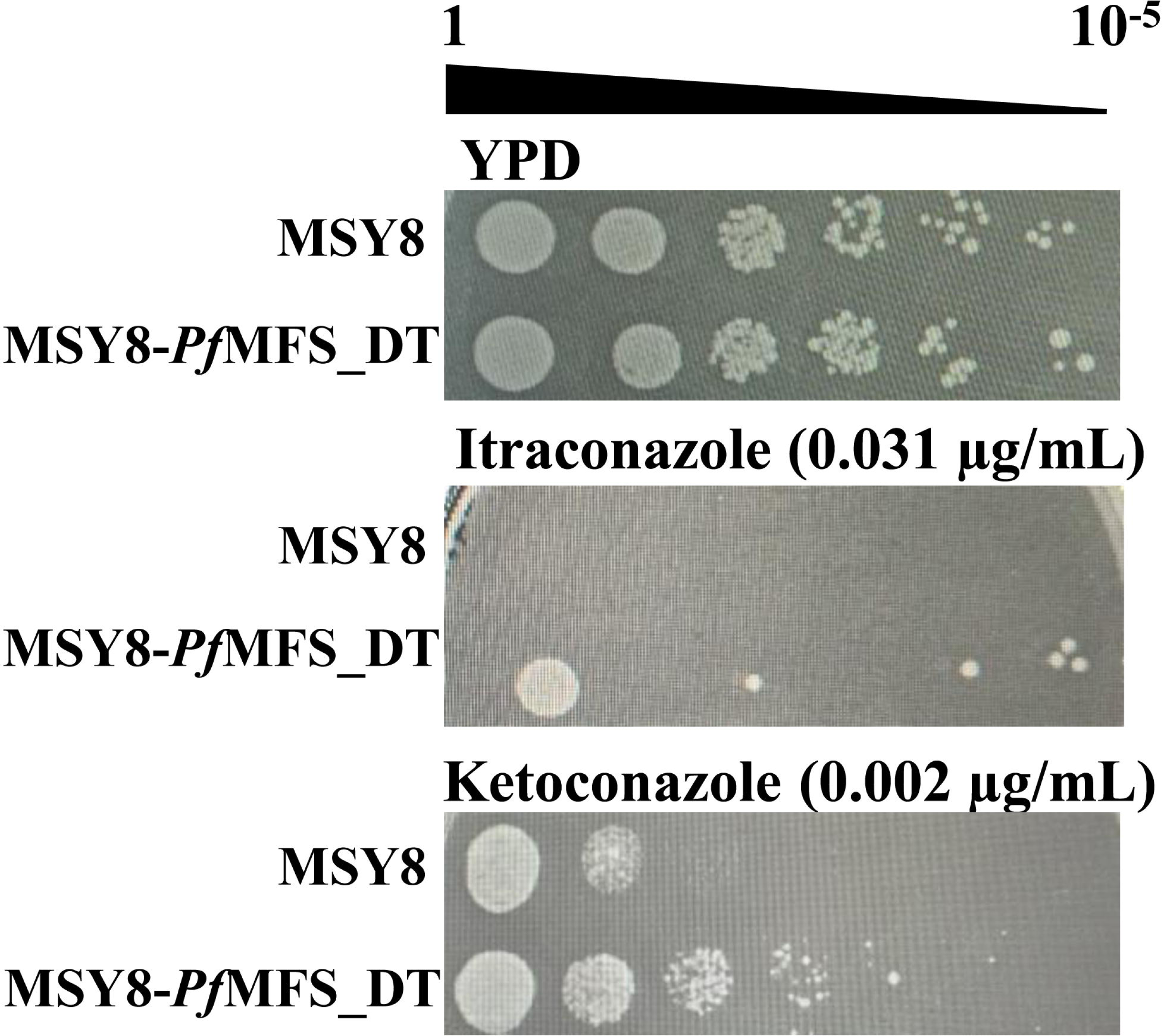
Drug susceptibility assay. Drug susceptibility assay was performed by serial dilution spot assay on transformed and untransformed MSY8, in the presence of itraconazole (0.031 µg/mL) and ketoconazole (0.002 µg/mL). *Pf*MFS_DT-expressing MSY8 cells showed rescued growth in the presence of itraconazole and ketoconazole.

### Antifungals, ketoconazole and itraconazole favorably interacts with *Pf*MFS_DT and inhibits *P. falciparum* growth

Homology modelling of *Pf*MFS_DT was done using Prime application in Schrodinger Maestro Suite 2022-4, using chain_A of bacterial oxalate transporter (OxlT 8HPK_A) [23] as a template. The generated model was further refined for its loop refinement and energy minimization using the OPLS4 force field in Schrodinger Prime [24] **(Figure 5A (i))**. The *Pf*MFS_DT model was mapped for its localization in the plasma membrane **(Figure 5A (ii))**. An analysis of the Ramachandran plot revealed that 94.9 % of the amino acid residues occupied the most favoured regions, with 4.6 % in the additional allowed regions **(Figure 5A (iii))**. A mere 0.2 % of the amino acids resided in generously allowed regions, while another 0.2 % in disallowed regions. These findings provide substantial support for the structural validity of the modelled structure, justifying its utilization in subsequent docking studies **(Figure 5A (iv))**.

**Figure 5:**
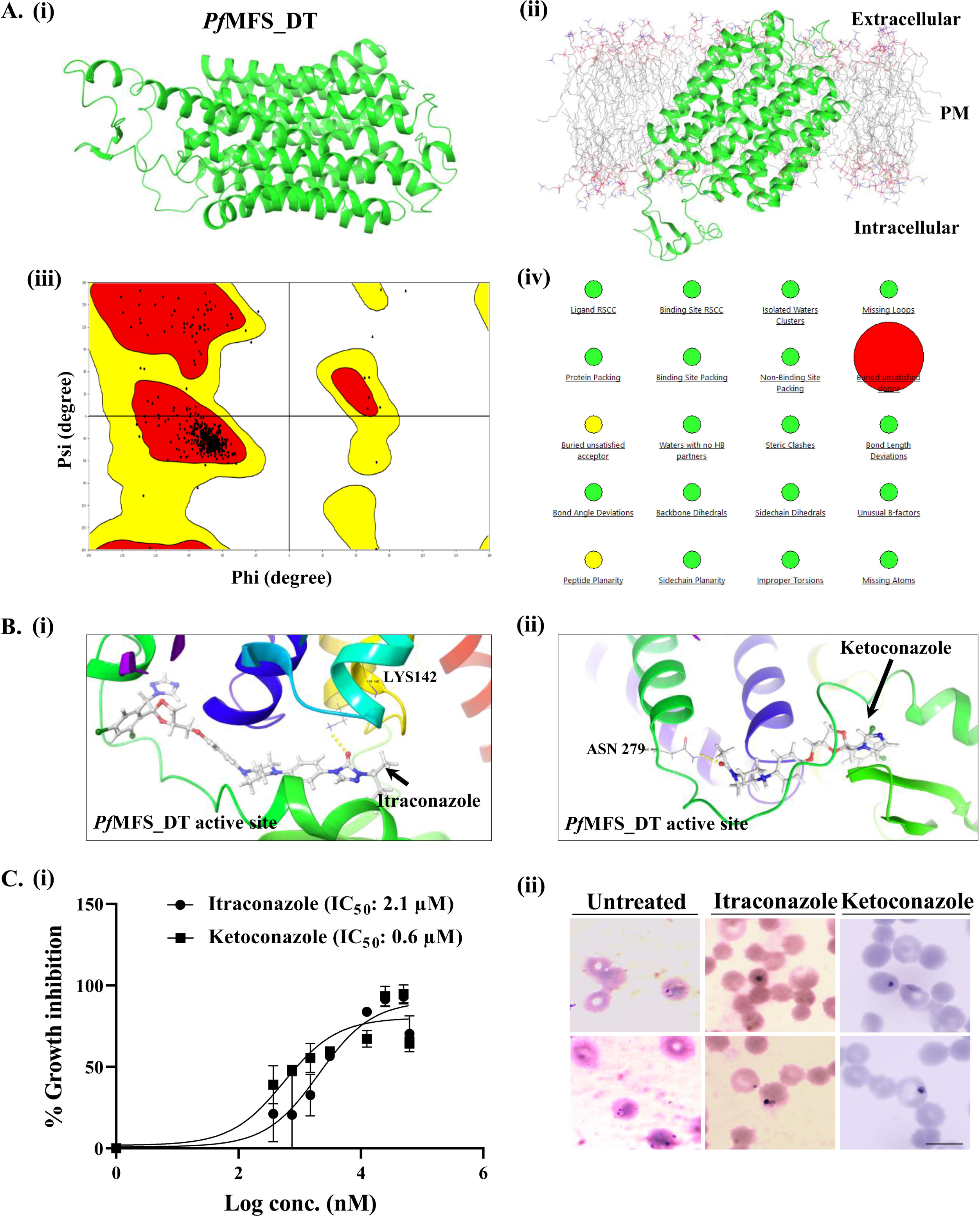
(A) Molecular modelling of *Pf*MFS_DT. Homology modelling of *Pf*MFS_DT was done using Prime application in Schrodinger Maestro Suite 2022-4, using chain_A of bacterial oxalate transporter (OxlT 8HPK_A) [23] as a template. The generated model was further refined for its loop refinement and energy minimization using the OPLS4 force field in Schrodinger Prime [24] **(i)**. *Pf*MFS_DT model was mapped for its localization in the plasma membrane **(ii)**. Ramachandran plot generated from PROCHECK [25] at the SAVES version 4 server for validation of generated *Pf*MFS_DT model **(iii)**. Plot analysis revealed 94.9% of the residues in the most favoured regions, 4.6% in the additional allowed regions, 0.2% of the amino acids in the generously allowed regions and 0.2% of the residues in the disallowed regions. Structural reliability of *Pf*MFS_DT **(iv)**. **(B) Molecular docking of itraconazole and ketoconazole.** Molecular docking of itraconazole **(i)** and ketoconazole **(ii)** against active site of *Pf*MFS_DT using Extra Precision (XP) docking [29–31]. Hydrogen-bonding, hydrophobic, and electrostatic interactions were assessed avoiding steric clashes. The Glide Score scoring function was applied to re-score the obtained poses. The Molecular Mechanics-Generalized Born Surface Area (MM-GBSA) method, utilizing the OPLS4 force field, VSGB solvent model, and search algorithms, was used [24], and the binding free energies of ligand-receptor complex were computed. The binding free energy for itraconazole is −64.91 kCal/mol whereas for ketoconazole is −73.8 kCal/mol. **(C) Antimalarial activity of itraconazole and ketoconazole.** *Pf*3D7 (ring stage, at 0.5% parasitemia) was treated with itraconazole and ketoconazole (0.37 to 50 µM) for 72 hours. Dose-response curve; IC_50_ values of itraconazole and ketoconazole were found to be 2.1 µM and 0.6 µM, respectively **(i)**. Giemsa-stained smears showing growth defects in drug-treated parasites as compared to the untreated control **(ii)**.

Ligands (ketoconazole and itraconazole) were docked against the active site of *Pf*MFS_DT using Glide [29] and OPLS-4 [24] force fields **(Figure 5B (i and ii))**. Glide module scores from XP docking within the predicted catalytic pocket of *Pf*MFS_DT were found to be - 1.599 and −4.519 for ketoconazole and itraconazole, respectively **(Table 1)**.

**Table 1:** Binding free energies of *Pf*MFS_DT complexation with ketoconazole and itraconazole.

| **Title** | **docking score** | **MMGBSA dG Bind (kCal/mol)** |
| --- | --- | --- |
| **ketocanozole-DB01026** | -1.599 | -73.8 |
| **Itraconazole-DB01167** | -4.519 | -64.91 |

Further, the intra-erythrocytic growth inhibition assay of *P. falciparum* strain 3D7 depicted effective inhibition of *Pf*3D7 growth in the nanomolar range, for both compounds **(Figure 5C (i and ii))**.

## DISCUSSION

Numerous independent investigations have indicated the importance of drug transporters belonging to the ATP-Binding Cassette (ABC) and Major Facilitator Superfamily (MFS) transporter superfamilies in conferring drug resistance. Consequently, an in-depth analysis of the functional roles played by these therapeutically relevant drug transporters becomes imperative. To address this, there is a continual need for an overexpression system that can serve as a platform to facilitate the comprehensive functional assessment of these transporters. In this context, our study delves into elucidating the functional role of the putative *P. falciparum* Monocarboxylate Transporter as a drug transporter by leveraging a genetically engineered strain of *Candida glabrata* (MSY8) as an orthologous expression system.

IFAs revealed *Pf*MFS_DT expression on the plasma membrane of mature trophozoites and schizonts of *P. falciparum* 3D7, further validated by Western blot analysis **(Figure 2)**. This spatial and temporal expression pattern of *Pf*MFS_DT indicates its potential functional role in parasite metabolism and its engagement in essential cellular processes, rendering it an appealing target for further research.

Due to the limited responsiveness of transporter proteins to proteomic tools, conducting proteomic and functional analysis on them can be challenging. Consequently, an alternative approach involves studying these transporters by expressing them in an orthologous system with a deleted background of major drug transporters. In this study, we utilized the *Candida glabrata* strain MSY8, generated by Sonam Kumari *et al.* [14], which has disruptions in seven clinically relevant membrane-localized ABC drug transporter genes: *CgSNQ2*, *CgAUS1*, *CgCDR1*, *CgPDH1*, *CgYCF1*, *CgYBT1*, and *CgYOR1*. Additionally, a gain-of-function mutation was introduced in the *CgPDR1* transcription factor, resulting in the hyperactivation of the *CgCDR1* locus [14]. Previous research has demonstrated the suitability of MSY8 for the functional analysis of non-ABC membrane transporters [14]. In this context, we orthologously expressed *Pf*MFS_DT in the *C. glabrata* MSY8 strain. This choice reduces the masking effects of dominant drug transporters on the activity of less-expressing transporters due to the deletion of endogenous transporters. Orthologous expression of *Pf*MFS_DT in MSY8 resulted in plasma membrane localization, as validated by Western blotting **(Figure 3)**, demonstrating the applicability of this yeast model for functional analysis of transporter proteins. The membrane localization of *Pf*MFS_DT in the yeast model suggested the conservation of its cellular targeting, further validating the MSY8 system for examining the transport activity of *Pf*MFS_DT.

The sensitivity of MSY8-*Pf*MFS_DT1 to ketoconazole and itraconazole, as demonstrated by spot assays, suggested the involvement of *Pf*MFS_DT in the transport of these antifungal drugs, emphasizing its potential role as a drug transporter **(Figure 4)**. This transporter-mediated antifungal transport was further supported by the rescued growth observed in the presence of ketoconazole and itraconazole. These antifungal drugs were used at their MIC_80_ concentrations; the lowest concentration of the test compounds inhibiting the growth of yeast by at least 80% relative to the drug-free YPD control is already known [14] for the MSY8 strain.

For further investigation, the homology modeling generated a stereochemically accurate structural model of *Pf*MFS_DT **(Figure 5A)**. The localization of *Pf*MFS_DT on the plasma membrane, as anticipated by the proposed structural model, reinforced its functional relevance. Given that transporters involved in nutrient metabolism can function as drug transporters [9], the sensitivity of MSY8-*Pf*MFS_DT1 towards antifungal drugs effective against malaria parasites, such as ketoconazole [32] and itraconazole [33], was evaluated. The *in silico* interaction analysis with antifungal compounds ketoconazole and itraconazole indicated favourable binding energies, implying probable binding interactions with *Pf*MFS_DT **(Figure 5B and Table 1)**. These findings enhance our understanding of the structural aspects of *Pf*MFS_DT and establish the foundation for further research into potential drug interactions and experimental validation of *Pf*MFS_DT as a target for antimalarial drug development. Furthermore, the nanomolar inhibitory effects of ketoconazole and itraconazole on intra-erythrocytic growth of *P. falciparum* strain 3D7 **(Figure 5C)** highlight their antimalarial properties, emphasizing the relevance of the yeast-based model system in identifying novel antimalarials.

*C. glabrata* strain MSY8-*Pf*MFS_DT1, which has been established in this study, offers a valuable tool for the screening of natural compounds, such as osthol, cumin, reserpine, caffeoylquinic acids, reserpine, piperines, and chalcones. These compounds, which are known to be inhibitors of multidrug transporters originating from plants [34,35]) may shed light on their potential interactions with *Pf*MFS_DT. Furthermore, an intriguing dimension to this study may be added by exploring whether *Pf*MFS_DT functions as an efflux pump for the conventional antimalarial drugs currently in use. Subsequent investigations are imperative to explore the precise mechanisms of action and evaluate the safety and efficacy profiles of these antifungal compounds *in vivo*.

## CONCLUSION

To understand the significance of *Pf*MFS_DT in drug transport, we adopted an integrative approach combining molecular biology, bioinformatics, and pharmacological assays. Towards this, we established a robust platform to orthologously express and investigate the functional significance of *Pf*MFS_DT by using *C. glabrata* MSY8 strain with disruptions in major drug transporter genes. The successful expression of *Pf*MFS_DT in MSY8, along with its susceptibility to the antifungal medications, ketoconazole and itraconazole, confirms its functional role as a drug transporter. Furthermore, we observed nanomolar inhibitory effects of the antifungals on the intra-erythrocytic proliferation of *P. falciparum*. Overall, our findings demonstrate the utility of the yeast-based model system (MSY8-*Pf*MFS_DT1) to identify novel *Pf*MFS_DT-targeting antimalarials, and propose *Pf*MFS_DT as a drug transporter.

## SUMMARY POINTS

**1.** In this study, we aimed to evaluate the functional relevance of a putative *P. falciparum* Major Facilitator Superfamily protein (*Pf*MFS_DT) in drug transport using genetically engineered *C. glabrata* lacking clinically relevant ABC drug transporter genes.
**2.** Our results indicate that *C. glabrata* strain MSY8 transformed with *Pf*MFS_DT conferred resistance to antifungal drugs, mirroring the resistance observed with another MFS transporter, *Cg*FLR1.
**3.** By combining bioinformatics, cellular and molecular biology, and pharmacological approaches, we have contributed to the understanding of *P. falciparum* physiology and the identification of a novel MFS protein as a drug transporter.

## Authors’ contributions

**Preeti Maurya:** Methodology, Validation, Formal analysis, Investigation, Data curation, Writing - Original Draft, and Writing - Review and Editing. **Mohit Kumar:** Methodology and Investigation. **Ravi Jain:** Methodology, Investigation, Writing - Original Draft, and Writing-Review and Editing. **Harshita Singh:** Methodology and Investigation. **Haider Thaer Abdulhameed Almuqdadi:** Methodology and Investigation. **Aashima Gupta:** Methodology and Investigation. **Christoph Arenz:** Writing - Review and Editing. **Naseem A. Gaur:** Resources, Writing - Original Draft, and Writing-Review and Editing. **Shailja Singh:** Conceptualization, Methodology, validation, Formal analysis, Resources, Data curation, Writing - Original Draft, Writing-Review and Editing, Supervision, Project administration, Funding acquisition.

## Acknowledgments

1. S. Singh is a recipient of the National Bioscientist award. PM is a recipient of Junior Research Fellowship from Department of Biotechnology, Government of India (Ref. no.: DBTHRDPMU/JRF/BET-21/I/2021-22/326). MK acknowledges Research Associate fellowship from Indian Council of Medical Research (ICMR-RA), Government of India (Myco/fell/3/2022-ECD-II). RJ acknowledges Research Associate I fellowship from Indo-DBT (Department of Biotechnology), Government of India and Swiss National Science foundation. AG acknowledges Laboratory Assistant fellowship from Indo-DBT (Department of Biotechnology), Government of India and Swiss National Science foundation.

## Writing disclosure

No writing assistance was utilized in the production of this manuscript.

## REFERENCES

1 World Health Organization (WHO). World Malaria Report 2023, (2023).

2 Counihan NA, Modak JK, de Koning-Ward TF. How Malaria Parasites Acquire Nutrients From Their Host. Front. Cell Dev. Biol. 9, 649184 (2021).

3 McKee RW, Ormsbee RA, Anfinsen CB, Geiman QM, Ball EG. The chemistry and metabolism of normal and parasitized (P. knowlesi) monkey blood. J. Exp. Med. 84(6), 569–582 (1946).

4 Kanaani J, Ginsburg H. Transport of lactate in Plasmodium falciparumLJinfected human erythrocytes. J. Cell. Physiol. 149(3), 469–476 (1991).

5 Pfaller MA, Krogstad DJ, Parquette AR, Nguyen-Dinh P. Plasmodium falciparum: Stage-specific lactate production in synchronized cultures. Exp. Parasitol. 54(3), 391–396 (1982).

6 Martin RE, Henry RI, Abbey JL, Clements JD, Kirk K. The ‘permeome’ of the malaria parasite: an overview of the membrane transport proteins of Plasmodium falciparum. Genome Biol. 6(3), R26 (2005).

7 Wunderlich J. Updated List of Transport Proteins in Plasmodium falciparum. Front. Cell. Infect. Microbiol. 12, 926541 (2022).

8 Kirk K. Channels and transporters as drug targets in the Plasmodium-infected erythrocyte. Acta Trop. 89(3), 285–298 (2004).

9. Liang Y, Li S, Chen L. The physiological role of drug transporters, (2015).

10 Piro F, Focaia R, Dou Z, Masci S, Smith D, Cristina M Di. An uninvited seat at the dinner table: How apicomplexan parasites scavenge nutrients from the host, (2021).

11 Beck JR, Ho CM. Transport mechanisms at the malaria parasite-host cell interface. PLoS Pathog. 17(4), e1009394 (2021).

12 Singh M, Afonso J, Sharma D et al. Targeting monocarboxylate transporters (MCTs) in cancer: How close are we to the clinics? Semin. Cancer Biol. 90, 1–14 (2023).

13 Sun Y, Sun J, He Z et al. Monocarboxylate Transporter 1 in Brain Diseases and Cancers. Curr. Drug Metab. 20(11), 855–866 (2019).

14 Kumari S, Kumar M, Khandelwal NK et al. A homologous overexpression system to study roles of drug transporters in Candida glabrata. FEMS Yeast Res. 20(4), foaa032 (2020).

15 Aurrecoechea C, Brestelli J, Brunk BP et al. PlasmoDB: A functional genomic database for malaria parasites. Nucleic Acids Res. 37(SUPPL. 1), D539–D543 (2009).

16 Omasits U, Ahrens CH, Müller S, Wollscheid B. Protter: Interactive protein feature visualization and integration with experimental proteomic data. Bioinformatics 30(6), 884–886 (2014).

17 Cater RJ, Chua GL, Erramilli SK et al. Structural basis of omega-3 fatty acid transport across the blood–brain barrier. Nature 595(7866), 315–319 (2021).

18 Liu W, Xie Y, Ma J et al. IBS: An illustrator for the presentation and visualization of biological sequences. Bioinformatics 31(20), 3359–3361 (2015).

19 Tamura K, Stecher G, Kumar S. MEGA11: Molecular Evolutionary Genetics Analysis Version 11. Mol. Biol. Evol. 38(7) (2021).

20 Corpet F. Multiple sequence alignment with hierarchical clustering. Nucleic Acids Res. 16(22), 10881–10890 (1988).

21 Trager W, Jensen JB. Human malaria parasites in continuous culture. Science (80-.). 193(4254), 673–675 (1976).

22 Bateman A. UniProt: A worldwide hub of protein knowledge. Nucleic Acids Res. 47(D1), D506–D515 (2019).

23 Jaunet-Lahary T, Shimamura T, Hayashi M et al. Structure and mechanism of oxalate transporter OxlT in an oxalate-degrading bacterium in the gut microbiota. Nat. Commun. 14(1), 1730 (2023).

24 Roos K, Wu C, Damm W et al. OPLS3e: Extending Force Field Coverage for Drug-Like Small Molecules. J. Chem. Theory Comput. 15(3), 1863–1874 (2019).

25 Laskowski RA, MacArthur MW, Thornton JM. PROCHECK: a program to check the stereochemical quality of protein structures. J. Appl. Crystallogr. 26(2), 283–291 (1993).

26 Jacobson MP, Pincus DL, Rapp CS et al. A Hierarchical Approach to All-Atom Protein Loop Prediction. Proteins Struct. Funct. Genet. 55(2), 351–367 (2004).

27 Sastry GM, Adzhigirey M, Day T, Annabhimoju R, Sherman W. Protein and ligand preparation: Parameters, protocols, and influence on virtual screening enrichments. J. Comput. Aided. Mol. Des. 27(3), 221–234 (2013).

28 Jacobson MP, Friesner RA, Xiang Z, Honig B. On the role of the crystal environment in determining protein side-chain conformations. J. Mol. Biol. 320(3), 597–608 (2002).

29 Halgren TA, Murphy RB, Friesner RA et al. Glide: A New Approach for Rapid, Accurate Docking and Scoring. 2. Enrichment Factors in Database Screening. J. Med. Chem. 47(7), 1750–1759 (2004).

30 Friesner RA, Banks JL, Murphy RB et al. Glide: A New Approach for Rapid, Accurate Docking and Scoring. 1. Method and Assessment of Docking Accuracy. J. Med. Chem. 47(7), 1739–1749 (2004).

31 Friesner RA, Murphy RB, Repasky MP et al. Extra precision glide: Docking and scoring incorporating a model of hydrophobic enclosure for protein-ligand complexes. J. Med. Chem. 49(21), 6177–96 (2006).

32 Pender GC, Ebong NO, Ebiladei L, Adikwu E. Antimalarial Assessment of Artesunate/Ketoconazole in a Mouse Model. Am. J. Biomed. Sci. Res. 19(4), AJBSR.MS.ID.002601 (2023).

33 Pongratz P, Kurth F, Ngoma GM, Basra A, Ramharter M. In vitro activity of antifungal drugs against Plasmodium falciparum field isolates. Wien. Klin. Wochenschr. 123(SUPPL. 1), 26–30 (2011).

34. Varela MF, Stephen J, Bharti D, Lekshmi M, Kumar S. Inhibition of Multidrug Efflux Pumps Belonging to the Major Facilitator Superfamily in Bacterial Pathogens, (2023).

35. Seukep AJ, Kuete V, Nahar L, Sarker SD, Guo M. Plant-derived secondary metabolites as the main source of efflux pump inhibitors and methods for identification, (2020).

