## Supplementary Material for "Expression of *Plasmodium* Major Facilitator Superfamily Protein in Transporters-Δ *Candida* Identifies a Drug Transporter"

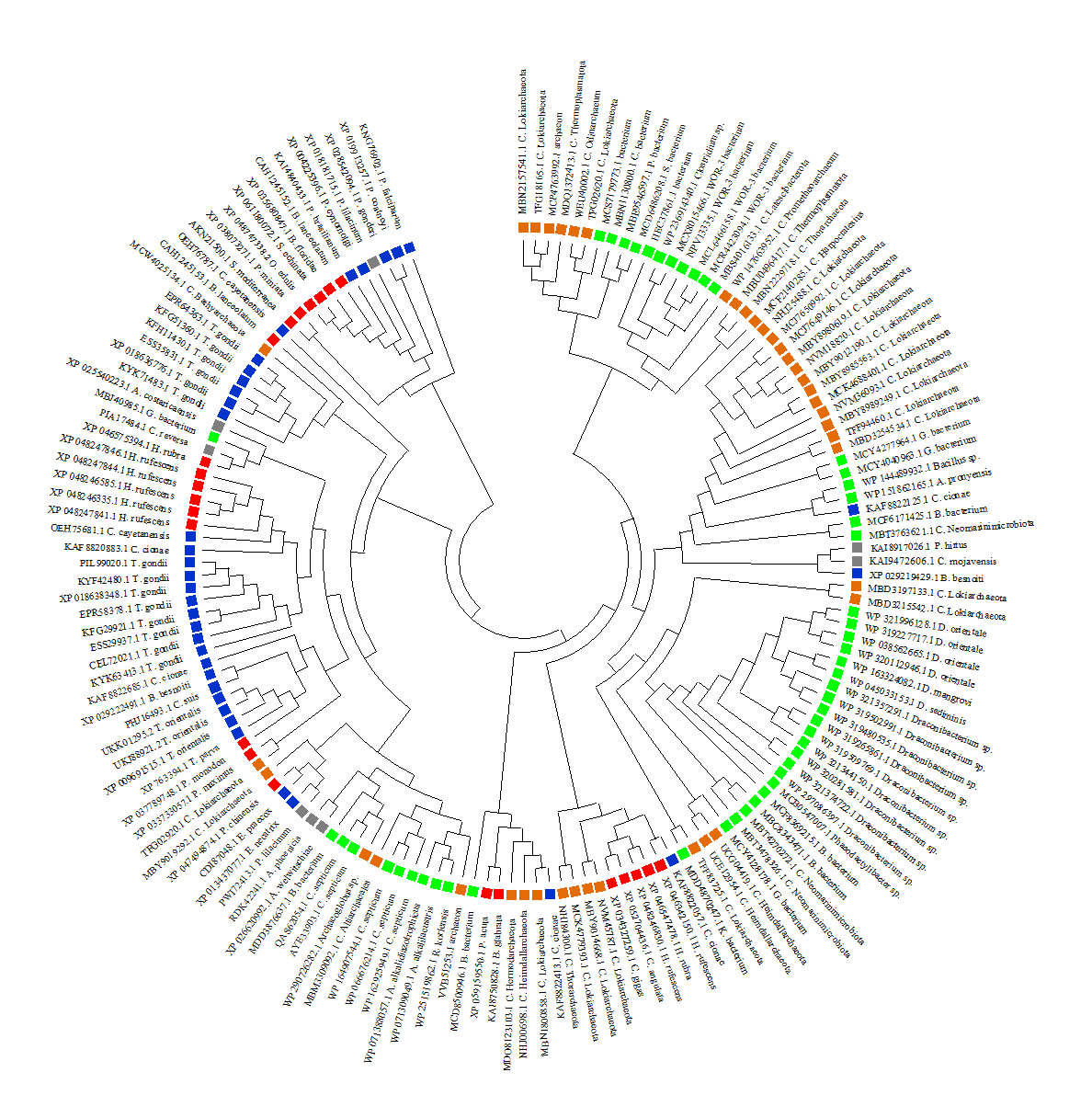

**Figure S1: The maximum likelihood phylogenetic relationship of *Pf*MFS-DT with its orthologs (MFS proteins) from various taxa, *viz*, archaebacteria, bacteria, apicomplexa, fungi, and animalia.** The evolutionary analysis was established using MEGA11, a user-friendly software suite for analyzing DNA and protein sequence data from species and populations (http://www.megasoftware.net/) (Tamura *et al.*, 2021). Branches with blue, brown, green, red, and grey represent MFS proteins from apicomplexans (taxid: 5794), archaebacteria (taxid: 2157), bacteria (taxid: 2), animalia (taxid: 33208), and fungi (taxid: 4751), respectively.

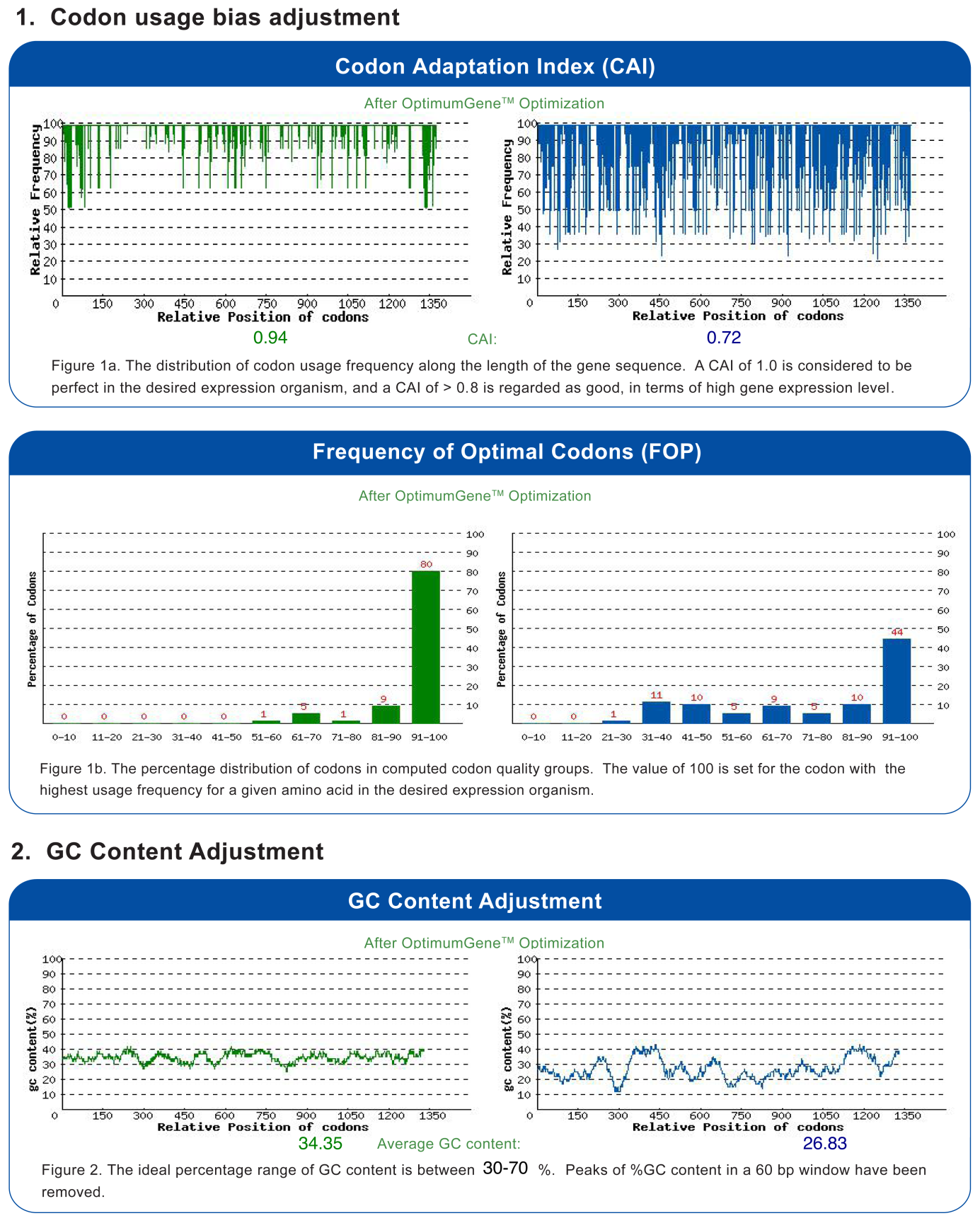

**Figure S2: Codon optimization report.** The Complementary Determining Sequence (CDS) encoding *Pf*MFS-DT (PlasmoDB ID: PF3D7_0210300) was codon-optimized for expression in yeast (GenScript Biotech, United States). Codon usage bias adjustment (panel # 1) and GC content adjustment (panel # 2) are shown.

**Original and codon-optimized nucleotide sequences of *Pf*MFS-DT:**

**> PF3D7_0210300_Original**

ATGAAAAAAGAGAATACTTCCCTGTTATCATCAAGATATAGAATAATATTAGGAGGATTTTTAATTCATTGTACGTTAGGAAGCATTTATTGTTTTTCTAATATAAGTGTATATGTAATATCATATATGAAAATAATAGGATGTTCTGATGTAAAATATAAAGATAGTAGTTGGATATATGTGTTGACTTTATTATTTCAATGTTTTTTTGGTTTTTTTGGAGGAATATTAAATCAGAATTTAGGACCACAGATTAGTGTCTTATTAGGTGGATGGTTAATGTGTTTAGGAATATTATTATCATATTTTACTGTTTTTAATTTTTATTTATTTTTAATGACTTATGGAATATTATGTGGAATAGGATGTGGAATAGCATATCCTATCCCTTTATCAGTGGCTGTCAAAAAGCACTATGATTACAAAGGAGTGATTAGCGGTATCATATTCATAGGGAGAGGACTTTCCGTGTTCATTATTTGCCCTTTACAGAATTATTACATAAATAAATATAATTATATGCCAGATTATATGCCCGAAATAGAAAACTCCGATGAGAAATATTTTAGTAACTTAGATATATTAAATAAGGTACCTTATTTGTTTATATATGAAGGAATATGTTTTGCTATTATTCAGTTTTTGGGTTCATATTTAATTGCAGATTCAGGTGATACATCTAAGGATTTCATGGCATATAATGATAGGAATAATAAAGTATTATATTTTGAAGAAAAAAATTTTATAAATAAGCCAAATGGTTTATCTAATTCTTTAAGAACATTATCGAATACATCGAATTTTTCATTTAGAGAAGTAAATAATACATTTATTAATCGTGAATTTATATTAATATGGTTAATGATCTTTTTTAATTGGCAAGCTATATCATATACTCAAGTATTTTGGAAAATATTTGGGATGAATTATTTATCTATTGATGATAGATCATTATCATTATTAGGATCTGTATCTTCTCTTTTTAATATTTTTGGTAGGATCTTTTGGGGACTTATAAGTGACTTTACAAGTTTTAAAACAACATTAATATTAATGAGTCTTCTTATGAGCTTTTTAACAATCACATTAACAATGTCAGGATTTTATGGTATTATAACTTATTCTATATGGGTATGTCTTATTTTCTTTTGTCATGCTGGCACTTTTGCAATATTCCCATCCATAACTGCCCATACATTTGGAACCAAAAATTTCGGACCAGTTTTTGGACTCCTATTTACAGCACGAGCTTTTTCAAGTATAATTAATGCAATCATATCAGCTGTCTTGTTAAATAATATTGGTAATATTGCAATGTGTGCAATTGTTTCCTTATCATCCTTTGTCAGCATCATGTTAGCACTAGCATTTTAA

**> PF3D7_0210300_Optimized**

ATGAAGAAAGAGAACACCAGTTTGCTTAGTAGTAGATACAGAATTATTTTGGGAGGATTTTTGATCCATTGCACA

TTGGGTAGTATTTACTGTTTCTCTAACATCTCTGTTTACGTTATTTCTTACATGAAGATCATCGGTTGTTCTGAT

GTTAAGTACAAGGATTCTTCTTGGATCTACGTTTTGACTTTGTTGTTCCAATGTTTCTTTGGTTTCTTTGGTGGT

ATTTTGAACCAAAATTTGGGTCCACAAATTTCTGTTTTGTTGGGTGGTTGGTTGATGTGTTTGGGTATTTTGTTG

TCTTACTTCACTGTTTTTAACTTCTACTTGTTTTTGATGACTTATGGTATTTTGTGTGGTATTGGTTGTGGTATT

GCTTATCCAATTCCTTTGTCTGTTGCTGTTAAGAAACATTACGATTACAAGGGTGTTATTTCTGGTATCATTTTC

ATTGGTAGAGGTTTGTCTGTTTTTATTATTTGTCCATTGCAAAACTACTACATCAACAAATACAACTACATGCCA

GATTACATGCCTGAAATCGAGAACTCTGATGAAAAGTACTTCTCTAATTTGGATATTTTGAACAAAGTTCCTTAC

TTGTTTATCTACGAGGGTATTTGTTTCGCTATCATCCAATTCTTGGGTTCTTATTTGATCGCTGATTCTGGAGAT

ACTTCTAAGGATTTCATGGCTTACAACGATAGAAACAACAAGGTTTTGTACTTCGAAGAGAAGAACTTCATCAAC

AAGCCTAACGGTTTGTCTAACTCTTTGAGAACTTTGTCTAACACTTCTAACTTCTCTTTCAGAGAAGTTAACAAC

ACTTTCATTAACAGAGAGTTTATTTTGATTTGGTTGATGATTTTCTTTAACTGGCAAGCTATTTCTTACACTCAA

GTTTTCTGGAAGATTTTCGGTATGAACTATTTGTCTATCGATGATAGATCCTTGTCTTTGTTGGGTTCTGTTTCT

TCTTTGTTTAACATCTTCGGTAGAATTTTCTGGGGTTTGATTTCTGATTTCACTTCTTTCAAAACTACTTTGATC

TTGATGTCTTTGTTGATGTCTTTCTTGACTATCACTTTGACTATGTCTGGTTTCTACGGTATCATCACTTACTCT

ATTTGGGTTTGTTTGATTTTCTTTTGTCATGCTGGTACTTTTGCTATTTTCCCATCTATTACTGCTCACACTTTC

GGTACTAAAAATTTTGGTCCTGTTTTCGGTTTGTTGTTTACTGCTAGAGCTTTTTCTTCTATTATTAACGCTATC

ATCTCTGCTGTTTTGTTGAACAACATTGGTAACATTGCTATGTGTGCTATCGTTAGTCTTTCCAGTTTTGTTTCC

ATTATGTTGGCATTGGCTTTC
